# Hijacking of internal calcium dynamics by intracellularly residing rhodopsins

**DOI:** 10.1101/2023.05.25.542240

**Authors:** Ana-Sofia Eria-Oliveira, Mathilde Folacci, Anne Amandine Chassot, Sandrine Fedou, Nadine Thézé, Dmitrii Zabelskii, Alexey Alekseev, Ernst Bamberg, Valentin Gordeliy, Guillaume Sandoz, Michel Vivaudou

## Abstract

Rhodopsins are ubiquitous light-driven membrane proteins with diverse functions, including ion transport. Widely distributed, they are also coded in the genomes of giant viruses infecting phytoplankton where their function is not settled. We examined the properties of three type 1 viral channelrhodopsins (VCR1s), and, unexpectedly, found that VCR1s accumulate exclusively intracellularly, and, upon illumination, induced calcium release from intracellular IP_3_-dependent stores. In vivo, this light-induced calcium release was sufficient to remote control muscle contraction and behavior in VCR1-expressing tadpoles. VCR1s are the first rhodopsins shown to natively confer light-induced Ca^2+^ release, suggesting an original mechanism for reshaping the response to light of virus-infected algae. The ability of VCR1s to photorelease calcium without altering plasma membrane electrical properties marks them as precursors for novel optogenetics tools, with potential applications in basic research and medicine.

## Introduction

Rhodopsins are ubiquitous integral membrane proteins found in many living organisms, from bacteria to man (*1*). Although their functions are diverse, they share the property of being driven by light, due to the presence within their structure of the light-isomerizable retinal (*2*). Many microbial rhodopsins are ion transporters, either photon-driven ion pumps like bacteriorhodopsins, or photon-gated ion channels like channelrhodopsins. When exogenously expressed in the plasma membrane of mammalian cells, channelrhodopsins provide a means to modify cell excitability with high spatiotemporal resolution upon illumination, a property at the origin of optogenetics.

Rhodopsin genes have been identified in the genomes of giant viruses (*3–5*). Metagenomic and phylogenetic sequence analysis shows that viral rhodopsins (VRs) are extremely abundant in marine environments. They form a monophyletic group of proteins within the rhodopsin superfamily that splits into two distinct branches: VRs of type 1 and type 2 (*3, 6–8*). VRs have the expected 7-transmembrane-helices topology of rhodopsins, but show only distant sequence similarity to microbial rhodopsins of known functions such as ChR2. The few VRs that have been functionally characterized in heterologous systems encode light-driven cation channels (Type 1 VirChR1 and VirRDTS) (*8*). These channelrhodopsins are thought to reside in the plasma membrane of the virus-infected phytoplankton host and to enhance phototaxis through a process linking plasma membrane ion flux to intracellular calcium which drives flagellar motion (*9–11*).

Here, we have studied 3 type 1 viral channelrhodopsins (VCR1s) having ∼60% aminoacid sequence similarity: OLPVR1 (Organic Lake phycodnavirus rhodopsin 1), VirChR1 and TARA150, both found by database mining and of unknown marine origin (*8*). In Xenopus oocytes and a mammalian cell line, we discovered that native OLPVR1 is strictly expressed intracellularly, localized to the endoplasmic reticulum, and triggers an increase in cytoplasmic calcium proportional to the light power applied. VirChR1 and TARA150 had similar phenotypes. Such function, so far unique among rhodopsins, uncovers an unexpected facet of the interactions of giant viruses with their phytoplankton hosts. It also suggests optogenetic use of VCR1s in a variety of cells where a straight link between intracellular calcium and cell function exists. As proof-of-concept of such use, we show here that light irradiation reversibly modified tail movements of OLPVR1-expressing frog tadpoles.

### OLPVR1 expression in oocytes elicits light-induced Ca^2+^-activated chloride currents

Whereas naïve oocytes produced no light-sensitive currents (Fig. 1*D*), illumination of oocytes expressing OLPVR1 induced a current which reached a peak within seconds, slowly desensitized during illumination, and rapidly disappeared in the dark. The amplitude of photocurrents correlated with the intensity of the light (Fig. 1*A&B*). The selectivity of OLPVR1 photocurrents was investigated by changing the ion composition of the bathing solution (Fig. 1*C&D*). Exchanging K^+^ for Na^+^ did not affect the amplitude of the currents. Lowering external Cl^-^ from 100 to 10 mM caused a drastic reduction in current amplitude while shifting the reversal potential from ∼-25 to ∼+10 mV (Fig. 1*D*). The chloride selectivity of OLPVR1 photocurrents and their strong outward rectification are properties of the endogenous Ca^2+^-activated Cl^-^ currents (CaCCs) of *Xenopus* oocytes, known to be carried by TMEM16A channels (*12, 13*). We, therefore, tested two TMEM16A inhibitors, Ani9 (*14*) and MONNA (*15*). Ani9 and MONNA, at a concentration of 30 μM in ND96 solution, blocked photocurrents by 93.4±1.4% and 98.9±0.4%, respectively (Fig. 1*E*). Full 100% inhibition was not achieved, more noticeably with Ani9, but this is more likely due to incomplete block of CaCC currents by these agents. Indeed, similar incomplete inhibition was obtained when Ani9 and MONNA were tested on CaCC currents induced by the release of internal Ca^2+^ subsequent to the activation of Gq-coupled M3 receptors (Fig. S1) (*16*).

**Fig. 1.**
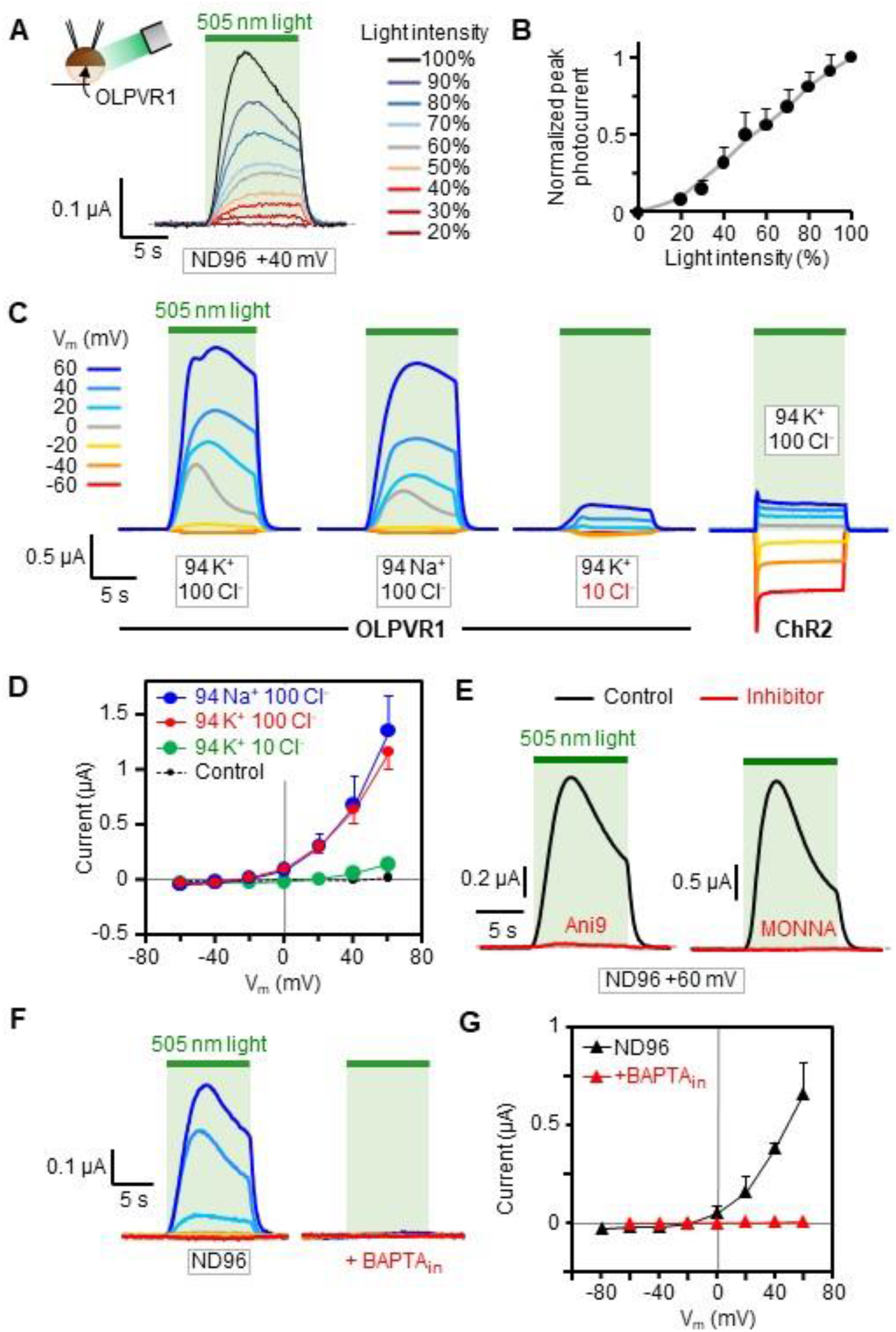
Photoactivation of OLPVR1 elicits CaCC currents in *Xenopus* oocytes. (A) Responses to a 10-s pulse of light of decreasing intensity. Current records were taken every 50 s from the same oocyte injected with 30 ng OLPVR1 RNA clamped at 40 mV. Bath solution was ND96. (B) Average peak current, normalized to current at 100% intensity, vs. light intensity obtained with the protocol of panel A, applied to 4 oocytes. (C) Photocurrents at different holding voltages from oocytes expressing OLPVR1 (30 ng RNA) or ChR2 (7.5 ng), in the specified different bath solutions. (D) OLPVR1 current-voltage relationships obtained from records as in panel C, in different extracellular ionic conditions. Control refers to currents recorded in non-injected oocytes. Currents were measured after 10-s illumination. Points are averages±SEM from 7-17 oocytes. (E) OLPVR1 photocurrents recorded before (Black) and after (Red) 60’ incubation in ND96 solution containing 30 μM Ani9 or MONNA, inhibitors of TMEM16A CaCCs. Statistics are shown in Fig. S1. (F) Photocurrents from OLPVR1-expressing oocytes (7.5 ng RNA) with and without intracellular injection of BAPTA (BAPTA_in_) in ND96 bath solution. (G) Average peak photocurrent vs voltage in ND96 solution with and without injected BAPTA. Each data point represents the average of 3-10 oocytes (Error bars, SEM).

Therefore, OLPVR1 expression reveals light-activated currents which have the features of the endogenous CaCC currents of oocytes.

### OLPVR1-mediated photocurrents require intracellular calcium

In order to understand the link between OLPVR1 and CaCC currents, we recorded photocurrents before and after microinjection of the fast Ca^2+^ chelator BAPTA in oocytes. Such treatment proved sufficient to eliminate all currents elicited by the activation of M3 receptors (Fig. S1). BAPTA injection in OLPVR1-expressing oocytes also eliminated all light-activated currents (Fig. 1*F&G*), demonstrating that photocurrents are mediated by changes in intracellular calcium. Whether oocytes were bathed in normal 1.8 mM extracellular Ca^2+^ (Fig. 1F&G), or elevated 49 mM Ca^2+^ (Fig. S2*B&D*), no photocurrent was detectable after BAPTA injection, given that the signal/noise ratio of our recordings afforded a resolution >∼1 nA. In particular, no inward current at negative potentials was observed, ruling out an influx of external Ca^2+^, directly or indirectly linked to OLPVR1.

We tested channelrhodopsin-2 (ChR2) as a control in the same conditions. As previously documented (*17*), the profile and BAPTA_in_ modulation of ChR2 photocurrents were completely different than those of OLPVR1 (Fig. S2*E-H*). In normal external Ca^2+^ (Fig. S2*E&G*), ChR2 cationic photocurrents were little affected by BAPTA_in_. In elevated external Ca^2+^ (Fig. S2*F&H*), ChR2, which is permeable to Ca^2+^ ions, produced currents different than in low Ca^2+^, especially at the most negative potentials where Ca^2+^ entry was large enough to activate CaCC currents (*17*). In agreement with this mechanism, BAPTA_in_ removed these large CaCC currents without affecting the intrinsic ChR2 currents.

Another remarkable difference between ChR2 and OLPVR1 is the significantly slower kinetics of the response to light of OLPVR1. The half-times of activation are ∼2 s for OLPVR1 and <0.1 s for ChR2 (Fig. S3). Fast activation of ChR2 photocurrents reflects the known intrinsic capabilities of ChR2 to sense light and conduct ions. Slow activation of OLPVR1 photocurrents is compatible with an action of OLPVR1 mediated by a diffusion-limited Ca^2+^-dependent process akin to the action of Gq-coupled M3 receptors (Fig. S3). In fact, CaCC response after OLPVR1 illumination was twice as slow as that after M3 activation by ACh.

These experiments show that, although both OLPVR1 and ChR2 can cause opening of CaCC channels, the mechanisms are fundamentally different. OLPVR1-induced photocurrents are entirely carried by CaCC channels activated by an elevation of intracellular Ca^2+^ independent of external Ca^2+^. ChR2 photocurrents result from cations flowing through the protein and, in conditions favoring Ca^2+^ permeation, an additional contribution from Ca^2+^-activated currents.

### Intracellular located OLPVR1 elicits release of calcium from IP_3_-dependent stores

Analysis of the atomic structure of OLPVR1 and its high phylogenetic similarity to VirChR1 (61% sequence identity) suggested that OLPVR1 could function like VirChR1 as a cation-conducting channel (*8*). The distinct effects of OLPVR1 in oocytes proved that it is expressed and functional. Our observations did not, however, show evidence of plasma membrane currents carried by OLPVR1 itself. This could be because OLPVR1 is not at the plasma membrane and we can only record the activity of channels present at the plasma membrane. To address the localization of OLPVR1, we used the XenoGlo technique, an adaptation of the Nano-Glo® system (Promega) to *Xenopus* oocytes. The HiBiT tag was inserted at the N-terminal extracellular end of OLPVR1 and ChR2, to produce functional OLPVR1_HB_ and ChR2_HB_ (Fig. S4). The data (Fig. 2A) show that OLPVR1 was not detected at the oocyte surface in contrast to ChR2 which was highly expressed.

**Fig. 2.**
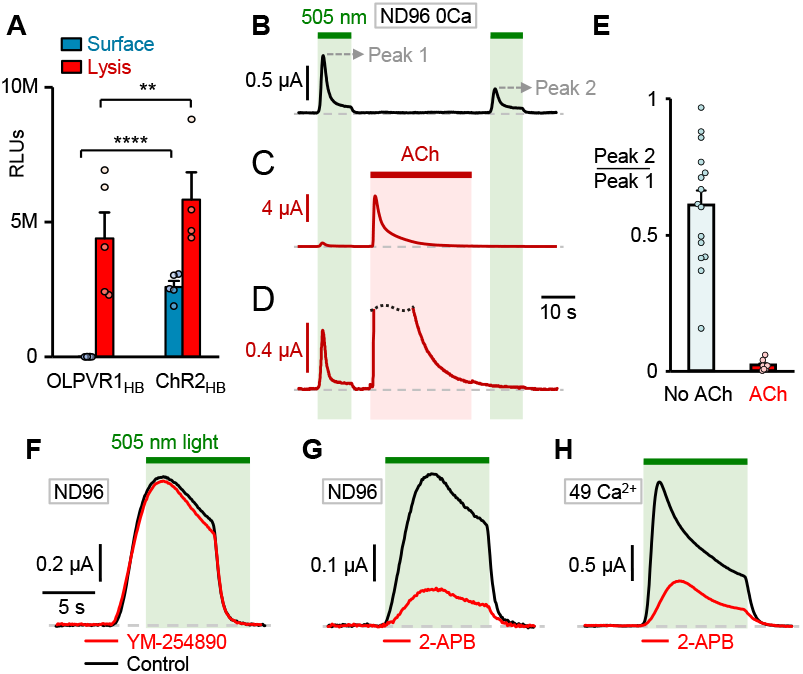
OLPVR1 is expressed intracellularly and activates surface CaCCs through release of intracellular Ca^2+^. (A) Surface expression of HiBit-tagged OLPVR1_HB_ compared to ChR2_HB_ measured using XenoGlo technique. OLPVR1, in contrast with ChR2, is not expressed as the surface membrane of oocytes. Mean luminescence recorded in oocytes injected with 7.5 ng RNA coding for OLPVR1_HB_ and ChR2_HB_ before (blue) and after (red) membrane permeabilization. The OLPVR1 surface value is 2360±700 RLUs. Values are from 4-7 batches of oocytes. (B) Photocurrents elicited by successive 10-s illuminations separated by a 40-s dark interval. Oocytes coexpressing OLPVR1 (30 ng RNA) and Gq-coupled muscarinic M3 receptor (2.5 ng) were bathed in ND96 0Ca solution. (C, D) Between the 2 illuminations, activation of M3 by ACh (5 μM; 30 s) induced Ca^2+^ release and large transient CaCC currents. Panel D is an enlarged version of C, showing a drastic reduction of the second photocurrent. (E) Average ratios of OLPVR1 peak current induced by the second illumination (Peak 2) over that of the first (Peak 1) with and without ACh application in between (Error bars, SEM). (F) Photocurrents at +60 mV in ND96 solution from oocytes expressing OLPVR1 (7.5 ng RNA) before (Control) and after 10’ incubation with 10 μM YM-254890.(G-H) Representative photocurrents at +60 mV in ND96 (G) or 49 Ca^2+^(H) solution from oocytes expressing OLPVR1 (7.5 ng RNA) before (Control) and after 60’ incubation with 100 μM 2-APB. Statistics are shown in Fig. S1.

Nonetheless, once oocytes were permeabilized, OLPVR1 was found in large amounts, comparable to the total amount of ChR2. This implies that OLPVR1 is not addressed to the surface membrane but is abundant in intracellular membranes.

As shown above, the intermediary between OLPVR1 photoactivation and CaCC currents is intracellular Ca^2+^. Could OLPVR1 modulate release of Ca^2+^ from intracellular stores? And, if so, which stores? These questions were addressed by observing the impact of Ca^2+^-stores depletion on OLPVR1 photocurrents. We coexpressed OLPVR1 and M3. Activation of M3 by its agonist Acetylcholine triggers the Gq-signaling cascade, the production of IP_3_, the release of Ca^2+^ stored in the endoplasmic reticulum (ER) by IP_3_ receptors, and the opening of CaCC channels (*16*). Under prolonged stimulation with a saturating ACh concentration (5 μM), in absence of extracellular Ca^2+^, the signature CaCC currents peaked within 2 s (Fig. 2*C*) and decayed by >99% within 30 s, an indication that Ca^2+^ stores have been emptied. We measured OLPVR1 photocurrents before and after such ACh application (Fig. 2C*-E*) and observed that photocurrents were barely detectable after ACh application, their amplitude decreasing to 2±0.6% of the initial amplitude. In control conditions, the same protocol without intervening ACh application showed a decrease of photocurrents of 39±6% (Fig. 2*B&E*).

These experiments suggest that the Ca^2+^ ions released by OLPVR1 come from the same stores as those released by IP_3_ receptors, presumably the endoplasmic reticulum. We also note that M3 stimulation produced ∼20-fold larger currents than OLPVR1. Such difference is not unexpected considering the different natures of the stimuli. Bath-applied ACh reaches the entire population of surface M3 receptors and the produced IP_3_ molecules can diffuse to reach most intracellular IP_3_ receptors. In contrast, light can only penetrate a few microns below the surface of opaque oocytes and reaches a small fraction (<5%) of the OLPVR1 proteins present (See Methods).

### OLPVR1 integrates tightly with the calcium-release mechanism of *Xenopus* oocytes

To further examine the link between Ca^2+^ release and OLPVR1, we utilized 2 inhibitors: YM-254890 targets Gq proteins (*18*), and 2-aminoethoxydiphenyl borate (2-APB) targets both IP_3_ receptors and Store-Operated Calcium Entry (SOCE) (*19*). Their effects on OLPVR1 photocurrents were tested at saturating concentrations that blocked >98% of the response to M3 activation in oocytes (Figs. 2*F-H* *& S1)*. The Gq inhibitor YM-254890 did not affect the OLPVR1 photocurrents, confirming that OLPVR1 mobilizes calcium release independently of the Gq pathway. 2-APB did not have such a clear-cut effect. It decreased OLPVR1 currents considerably but far from completely. On average, 2-APB reduced photocurrents amplitude by 75%. This suggests that the OLPVR1 photocurrents are largely contributed by the action of IP_3_R and/or SOCE, likely as a consequence of the positive feedback amplification of calcium signaling. This observation applies with the caveat that we do not know the direct effects of 2-APB on OLPVR1, which might confound our conclusions.

The fact that 2-APB fully blocks IP_3_-dependent currents elicited by M3 activation, but only partially block OLPVR1-induced photocurrents, suggests that OLPVR1 can induce a release of Ca^2+^ from stores even when IP_3_ receptors are inoperative. Thus, OLPVR1 can act independently of IP_3_ receptors although it appears that IP_3_ receptors contribute a large fraction of the OLPVR1 photocurrents – the 2-APB-sensitive fraction – likely because the Ca^2+^ release initiated by the fraction of illuminated OLPVR1s is propagated through opening of Ca^2+^-activated IP_3_ receptors (*20, 21*).

### ER-localized OLPVR1 increases intracellular calcium in HEK293T cells

To generalize our results, we turned to a mammalian cell line, HEK293T cells, and recorded currents with the patch clamp technique in the whole-cell configuration. In that configuration, the cytoplasm is perfused with the pipette solution. Unless otherwise specified, we added 10 μM of the slow chelator EGTA, a quantity sufficient to chelate the contaminant Ca^2+^ in our pipette solutions, but low enough to preserve Ca^2+^ signals as demonstrated below.

Initial experiments showed that light induced no detectable current in HEK293T cells transfected with OLPVR1 (Fig. 3*B &* Fig. S5). Accordingly, confocal fluorescence imaging showed that OLPVR1 was not expressed at the plasma membrane but co-localized with an ER marker (Fig. 3*A*). Unlike *Xenopus* oocytes, HEK293T cells lack endogenous Ca^2+^-sensitive channels to serve as electrophysiological readouts of intracellular Ca^2+^. To match oocyte conditions, HEK293T cells were transfected with OLPVR1 and TMEM16A (aka ANO1) – a gene coding for the same Ca^2+^-activated chloride channels as the CaCCs of *Xenopus* oocytes (*12*). As shown in Fig. 3*C*, light pulses transiently induced outwardly rectifying currents reversing at ∼0 mV, the expected chloride Nernst potential. Similar results were obtained with TMEM16B/ANO2, a homologue of TMEM16A found in neuronal cells (*22*) (Fig. S5).

**Fig. 3.**
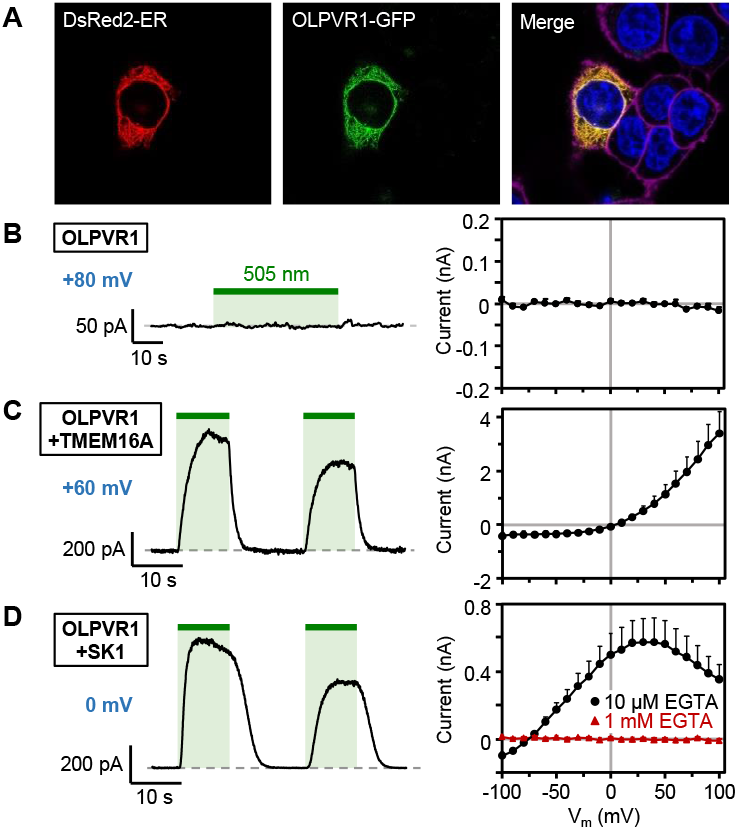
In mammalian cells, OLPVR1 localizes to the ER and activates surface Ca^2+^-activated channels through release of intracellular Ca^2+^. (A) Confocal images of HEK293T cells cotransfected with plasmids for expression of the ER-marker DsRed2-ER (red) and GFP-fused OLPVR1 (green). Merge panel shows in yellow, overlapping ER (red) and OLPVR1 (green) signals. The nucleus is in blue and the plasma membrane in magenta. (B-D) Whole-cell recordings were obtained from HEK293T cells transfected with OLPVR1 alone (A) or with OLPVR1 and, either TMEM16A (C) or SK1 (D). Left panels : Current responses to illumination (green bars, 505 nm light). Cells were held at the indicated voltages. Right panels: Light-induced current (peak current elicited by first illumination – current before illumination) vs. voltage. Currents were elicited by 400-ms voltage ramps from -100 to +100 mV repeated every second. Averages were calculated from 8 cells in each condition (Error bars, SEM). The pipette solution had 10 μM EGTA, except for the current-voltage curve drawn in red (right panel of D) where it had 1 mM EGTA.

Further tests were performed with a different Ca^2+^ sensor, the small-conductance Ca^2+^-activated potassium channel SK1 (*23*). Cells transfected with OLPVR1 and SK1 displayed large photocurrents that reversed near -80 mV, the K^+^ Nernst potential, and had a typical SK1 current-voltage relationship (Fig. 3*D*). The photocurrents disappeared when the slow chelator EGTA was included in the pipette solution at 1 mM, ruling out the type of tight association between the OLPVR1-dependent Ca^2+^ source and surface channels that has been found between IP_3_R and CaCCs (*24*).

### Other viral rhodopsins

OLPVR1 belongs to viral rhodopsin group 1. We tested two proteins from the same group, identified in Tara Ocean Foundation marine genomic data (*8, 25*): VirChR1 and TARA150. As shown in Fig. 4A&B, TARA150 and VirChR1 generated photocurrents that resembled the CaCCs currents generated by OLPVR1. The tested constructs were modified by addition of SS and/or MT sequences at their termini (See Methods) as these gave more robust responses. Such modifications, although supposed to promote surface expression, did not change the expression profile when used with OLPVR1 (Fig. S6). We further tested VirChR1 in HEK293T and observed light responses akin to those seen with OPLVR1, i.e, absence of photocurrents in VirChR1-transfected cells and large photocurrents in cells co-transfected with TMEM16A (Fig. 4C&D). In these experiments we used the same modified protein (HA-VirChR1-MT) as in a previous study using SH-SY5Y human neuroblastoma cells (*8*). In those cells, VirChR1, but not OLPVR1, produced detectable whole-cell currents indicative of a Na^+^/K^+^ permeable channel downregulated by external Ca^2+^ higher than ∼1 mM. Because the pipette contained 10 mM EGTA, any process involving intracellular calcium was suppressed.

**Fig. 4.**
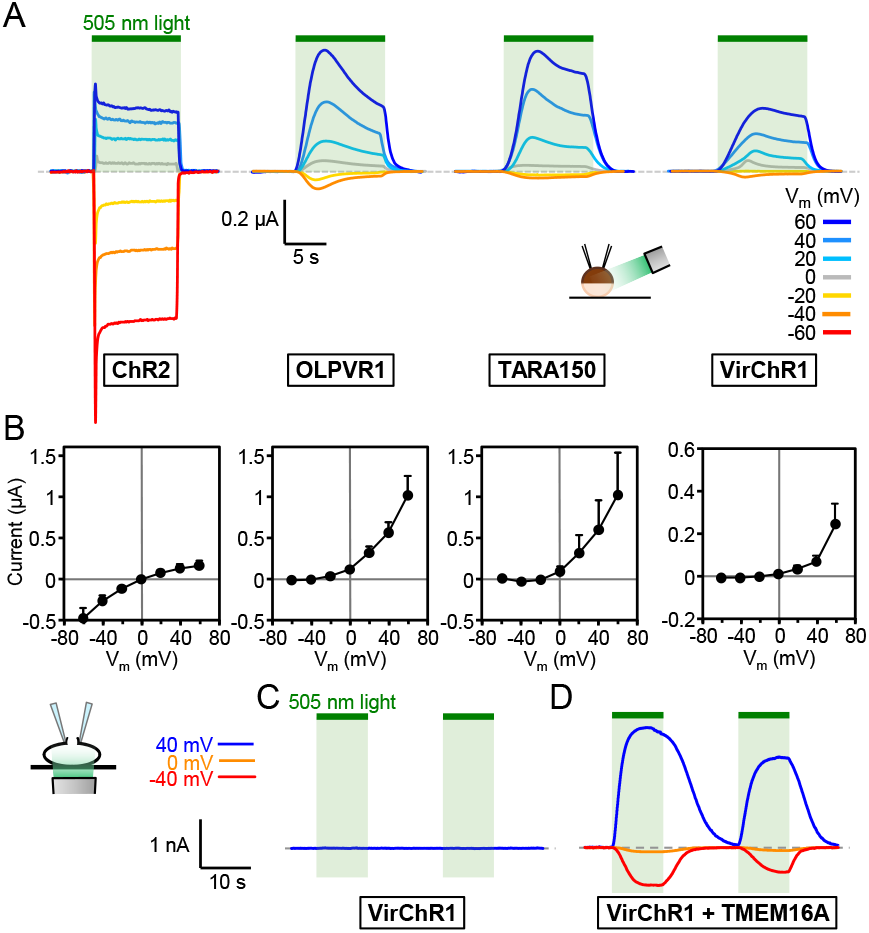
Photo-induced currents of members of the VR1 family have a similar profile which is distinct from that of ChR2. (A) Representative recordings in 94 K^+^ 100 Cl^-^ solution from oocytes expressing, from left to right, ChR2 (7.5 ng RNA), and the viral rhodopsins OLPVR1, TARA150 (construct SS-TARA150), and VirChR1 (construct SS-VirChR1-MT) (30 ng). (B) Current-voltage relationships of currents measured after 10 seconds of illumination, in KCl solution. Averages are from 3-10 oocytes. (C) Representative whole-cell patch clamp recordings from a HEK293T transfected with HA-VirChR1-MT. Standard bath and pipette solutions. (D) *Idem* from a cell co-transfected with VirChR1 (construct HA-VirChR1-MT) and TMEM16A.

### Light-driven animal motion endowed by OLPVR1

Calcium release is the trigger of muscle contraction. To test whether OLPVR1 can be expressed in living animals and produce light-dependent behavioral changes, we expressed OLPVR1 in amphibian *Xenopus laevis* tadpoles. Expression was achieved by injection of 1 ng OLPVR1 mRNA in one cell of two-cell stage *Xenopus* embryos. Testing was performed within 4 days of development after embryos had developed into free-swimming tadpoles. Results are illustrated in Fig. 5 (see also supplementary videos). Illumination with 505-530 nm light produced distinct motion in almost half of OLPVR1-injected tadpoles while it had no effect in control tadpoles injected with ß-Gal. We did not quantify expression of OLPVR1 but we know from experiments with single oocytes that protein expression after mRNA microinjection is highly variable. The responses were a combination of tail flicking, swimming and twitching (supplementary videos 4, 5 and 6, respectively). Tail flicking started and ended within 2 s when light was switched on and off. Swimming started more slowly, within 6 s, and stopped after >15 s. Such behaviors are consistent with light-induced Ca^2+^ release and muscle contraction. We however cannot rule out, at this juncture, OLPVR1 effects on other tissues, especially neuronal cells, since expression was not targeted to any particular cell type.

**Fig. 5.**
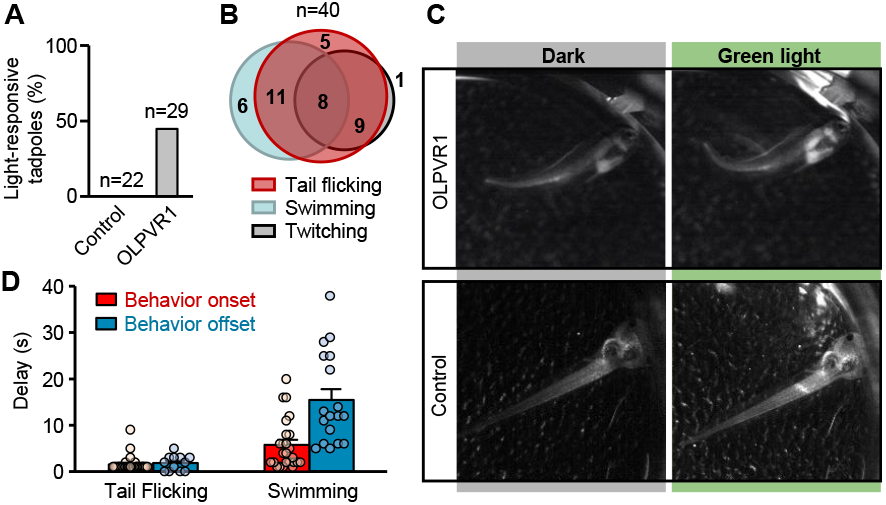
Green light induces motion of *Xenopus* tadpoles expressing OLPVR1. (A) Tadpoles from 2 independent batches of mRNA-injected embryos (Control injected with 250 pg lacZ mRNA, and OLPVR1 injected with 1 ng OLPVR1 mRNA) were subjected to 2-200-s green-light pulses and observed for light-triggered behavior. No lacZ tadpole (n=22) demonstrated green-light response. 44.8% of OLPVR1 tadpoles (n=29) displayed green-light-dependent motion. (B) Light-induced responses of a single OLPVR1-expressing tadpole (n=40 illuminations) were variable and consisted of a combination of tail flicking as in (C), swimming or post-illumination body twitching (see supplementary videos). (C) Images of tadpoles 3 days after injection of embryos with LacZ (Control) or OLPVR1 mRNA before (Dark) and after a 10-s illumination. (D) Delays of onset after light application, and offset after light switch-off, of behavioral response of OLPVR1 tadpoles. Tail flicking had onset and offset of 1.5±0.3 and 1.8±0.5 s, respectively. Swimming had onset and offset of 5.7±1.1 and 15.4±2.3 s, respectively. (Error bars, SEM)

## Discussion

Evidence in two different cell types demonstrates that native OLPVR1 localizes intracellularly in the ER and, upon light irradiation, mediates a rise in intracellular Ca^2+^ through release from IP_3_-dependent Ca^2+^ stores. Such so far unreported phenotype appears not to be an isolated outlier as it was shared by the two other related VCR1s, VirChR1 and TARA150, that were tested here.

How do VCR1s increase cytoplasmic Ca^2+^? The most parsimonious mechanism would be for these proteins to serve as a Ca^2+^-release channel in the ER membrane. In oocytes and HEK293T cells, whole-cell currents could not be obtained to investigate ion selectivity. The only information is from neuroblastoma cells (*8*) where VirChR1 expressed weakly, but sufficiently, at the surface membrane to show a Na^+^/K^+^ permeable channel blocked by extracellular Ca^2+^ above ∼2 mM. This block by Ca^2+^ is significant because it ensures that, in the high-Ca^2+^ environments of algal hosts (seawater, hypersaline Organic Lake), any VR present at the outer surface membrane of algae would be inactive. For ER-localized VCR1s, this block would be negligible because the external face of the channel would face the ER lumen where Ca^2+^ concentration does not exceed hundreds of μM (*26*). The ER membrane incorporates a variety of channels permeable to monovalent ions (*27*) which are thought to provide counterions currents to balance Ca^2+^ fluxes (*28*). In that context, increasing the permeability of the ER membrane to monovalent cations by opening VCR1s might facilitate Ca^2+^ release through ER Ca^2+^-leak channels (*29*). Many rhodopsins are proton pumps and alkaline pH in the ER lumen inhibits Ca^2+^ uptake by the Ca^2+^-ATPase pump, thus promoting an increase in cytosolic Ca^2+^ (*30*). OLPVR1 can also pump protons, but very weakly and in a direction that would instead acidify the ER lumen (*8*). Besides monovalent cations and protons, calcium ions could also permeate the VCR1s but at such low rates that they would not produce a measurable current and would appear to block the permeation of other ions. This mechanism would resemble the leaking of Ca^2+^ ions through voltage-dependent Na^+^ channels responsible for transient intracellular Ca^2+^ rise in neuronal axons (*31*).

If animal rhodopsins natively reside in internal membranes (*32*), it is accepted that microbial channelrhodopsins, such as *Chlamydomonas reinhardtii* ChR2 (*33*), natively localize and function at the surface membrane. Our results challenge this view by demonstrating that channelrhodopsins can have a biological function in internal membranes. When expressed in heterologous systems, microbial rhodopsins often accumulate in intracellular domains and have to be subjected to optimizations for surface membrane expression (*34*). In *Xenopus* oocytes, we found that a sizeable fraction of ChR2 proteins is intracellular but illumination of ChR2-expressing oocytes did not elevate intracellular Ca^2+^, at least to levels sufficient to activate CaCCs. This suggests that internal ChR2, unlike VCR1s, is not associated with Ca^2+^ stores. In contrast, when ChR2 was engineered to target the ER, its photoactivation caused elevation of intracellular Ca^2+^ (*35*). Precise targeting of VCR1s to the ER appears to be the key factor in their ability to trigger Ca^2+^ release and fulfill their cellular function. Preferential localizations cannot be explained by already described retention or export signal sequences. The amino acid sequences of VCR1s when subjected to localization prediction tools (*36*) showed no consistent result.

Organic Lake *phycodnavirus* is thought to infect prasinophytes or prymnesiophytes, members of unicellular algal flagellates, the most probable host being the prasinophyte *Pyramimonas* (*37*). The viruses carrying VirChR1 and TARA150 are unidentified. Channelrhodopsins in better known algal organisms reside in the plasma membrane at the eyespot and are the precursor of a phototransduction cascade involving depolarization, voltage-dependent Ca^2+^ channels activation, Ca^2+^-induced Ca^2+^ release from intracellular stores, and eventually flagellar motion and phototaxis (*9–11, 38*). Such a scheme would be altered in infected algae, with VCR1s providing a direct link between irradiation and intracellular Ca^2+^, bypassing any electrical signalling. VCR1s could modify the behaviour of their hosts to favour virus replication, possibly conferring or enhancing phototaxis (*4*). Hijacking intracellular Ca^2+^ dynamics is a strategy of many viruses. Notably, some Ca^2+^-permeable viroporins (virus-encoded ion channels) are targeted to the host ER membrane and modulate intracellular Ca^2+^ homeostasis to promote or prevent host apoptosis (*39, 40*). By analogy, VCR1s could also confer light-dependent protection against, or susceptibility to, apoptosis.

Ultimately, VCR1s connect the fundamental energy source of the host – light – to the most ubiquitous intracellular messenger – calcium. As such, they are expected to be key players in how giant viruses modulate population dynamics of their phytoplankton hosts and impact global ecology.

The precise release of calcium from intracellular stores mediates a panoply of transduction cascades incellular processes and pathologies such as gene expression, neurotransmitter release, hormone release, muscle contraction, host-pathogen interactions, tumour development or neurodegenerative disorders. Mobilization of intracellular calcium is achievable by pharmacological approaches (*21*). However, leveraging the spatiotemporal resolution of light delivery could enable more precise control of calcium release with potential advantages to our understanding of calcium signaling. The propensity of VCR1s to accumulate in internal storages with little or no sequence engineering and modulate calcium release, in a tunable manner depending on the light intensity, makes them attractive optogenetic tools, with potential applications in the manipulation of numerous aspects of cell activity and, as demonstrated, of animal behavior.

## Acknowledgments

We thank Pierre Thiébaud (Univ. Bordeaux, France) for assistance with Xenopus tadpoles, Jean Revilloud (IBS, Grenoble, France) for technical assistance, Christophe Moreau and Justine Magnat (IBS) for Xenopus frogs maintenance and oocytes preparation, Bruno Allard and Vincent Jacquemond (Univ. Lyon, France) and Maximilien Fürthauer (iBV, Nice, France) for preliminary tests, Christoph Fahlke (Inst. of Biological Information Processing, Jülich, Germany) for ChR2. Microscopy was performed in the MICA facility of the iBV-CNRS UMR 7277-INSERM U1091-UNS and we acknowledge the help of PRISM engineers Sameh Ben-Aicha and Baptiste Monterosso.

## Funding

Agence Nationale de la Recherche grant ANR-15-CE11–0029–01 (MV, VG)

Agence Nationale de la Recherche grant ANR-19-CE11-0026 (MV, VG, GS)

Centre National de la Recherche Scientifique CNRS ((MV, VG, GS)

Commissariat à l’Energie Atomique et aux Energies Alternatives CEA (MV, VG)

Université Grenoble Alpes (MV, VG)

CEA-IBS and Research Center Jülich FZJ-IBI7 collaborative agreement (VG)

French National Laboratories of Excellence “Ion Channel Science and Therapeutics” (LabEX ICST) network grant ANR-11-LABX-0015-01 (MV, GS)

Agence Nationale de la Recherche doctoral fellowship (ASEO)

Commissariat à l’Energie Atomique et aux Energies Alternatives CEA doctoral fellowship (MF)

## Author contributions

Conceptualization: ASEO, MV, VG

Formal analysis: ASEO, MV, DZ, AA

Methodology: ASEO, MF, MV, GS

Software: MV

Resources: DZ, AA, EB, SF, NT

Investigation: ASEO, AAC, MF, SF, NT

Visualization: ASEO, MV, AAC, GS

Funding acquisition: MV, GS, VG, EB

Project administration: MV, GS, VG

Supervision: MV, GS, VG, EB

Writing – original draft: MV, ASEO

Writing – review & editing: MV, ASEO, GS, DZ, VG, AA, AAC, MF, EB, SF, NT

## Data and materials availability

All data are available in the main text or the supplementary materials.

## Supplementary Materials

### Materials and Methods

#### Molecular biology

For experiments in *Xenopus* oocytes and tadpoles the genes of OLPVR1 (GenBank: ADX06642), OLPVR2 (GenBank: ADX06595), TARA150 (GenBank: MAV65030 with additional N-terminal sequence MVGGSL), VirChR1 (TARA-146-SRF-0.22-3-C376786_1), and ChR2 (fused to mKate red fluorescent protein; kindly provided by Prof. Christoph Fahlke, Jülich, Germany) were subcloned in the pXOOM vector (*41*), whilst the human muscarinic receptor M3 (GenBank: NP_001362914) and β-gal were subcloned in a pGEMHE vector (*42*). Protein constructs OLPVR1_HB_ and ChR2_HB_ were fused at the extracellular N-terminal with the HiBiT tag (sequence: VSGWRLFKKIS) followed by a GSSGGS linker. For that purpose, the corresponding nucleotidic sequences were inserted by standard PCR after the start codon of the OLPVR1 and ChR2 genes. mRNA coding for all proteins was prepared *in vitro* using the mMessage mMACHINE T7 Transcription Kit (#AM1344, Invitrogen) and purified using the NucleoSpin RNA XS purification kit (#740902, Macherey-Nagel).

For experiments in HEK293T (#CRL11268, ATTC) the genes were subloned as follows: OLPVR1 in pIRES2-EGFP, mouse TMEM16A (NP_848757), TMEM16B (NP_705817) in pmCherry-N1, and human SK1 (isoform 1, NP_001373903) in pcDNA3.1.

Where noted, sequences were added at the N-terminus (HA, hemagglutinin epitope; SS, N-ter signal sequence of 25 amino acids from human nicotinic acetylcholine alpha 7 receptor subunit isoform 1 (*43*) and the C-terminus (MT, C-ter Golgi export trafficking signal sequence – KSRITSEGEYIPLDQIDINV – of Kir2.1 channels (*44, 45*).

#### *Xenopus* oocytes

*Xenopus* oocytes were prepared as previously described (*46, 47*). Animal handling and experiments fully conformed with European regulations and were approved by the Ethics Committee of the *Commissariat à l’Energie Atomique et aux Energies Alternatives* (Ethics Approval #12-040). Authorization of the animal facility has been delivered by the regional administration (*Préfet de l’Isère, authorization # D 38 185 10 001*).

Each oocyte was injected with 50 nl of water containing the indicated amount of mRNA in ng. Oocytes were injected with 2.5 ng of M3 mRNA, 7.5 ng of ChR2/ChR2_HB_ mRNA, and/or 7.5 to 30 ng of OLPVR1/OLPVR1_HB_, TARA150, VirChR1 mRNA (Lower, as well as higher, amounts yielded smaller currents). Microinjected oocytes were incubated in the dark at 19°C in modified Barth’s solution (in mM: 1 KCl, 0.82 MgSO_4_, 88 NaCl, 2.4 NaHCO_3_, 0.41 CaCl_2_, 0.3 Ca(NO_3_)_2_, 16 HEPES, pH 7.4) supplemented with 100 U/ml penicillin and streptomycin, 0.1 mg/ml gentamycin and 1 μM all-trans retinal (#R2500, Sigma-Aldrich). The oocytes were left to express proteins for 1 to 4 days before electrophysiological recordings were performed. Expression levels of ion channels in oocytes are notoriously variable, but less so when oocytes are from the same batch. Although we present averages from oocytes of different batches, we ascertained that our observations were valid when comparing oocytes from the same batches.

#### XenoGlo: Surface expression measurements in oocytes

Briefly, this method is based on the formation of an active Nanoluciferase by the high-affinity complementation of its two parts, a short 11-aminoacid peptide (HiBiT) inserted as as a tag in the target protein, and a larger soluble LgBiT protein.

Oocytes injected with mRNA coding for HiBiT-tagged proteins such as OLPVR1_HB_ and ChR2_HB_ were dispensed with the animal pole facing up in white round-bottom 96-well plates pre-filled with 100 μl of modified Barth’s solution in each well.

Surface luminescence measurements of oocytes were performed using the kit Nano-Glo HiBiT Extracellular Detection System (#N2421, Promega) by adding to each well 100 μl of the Nano-Glo HiBiT extracellular reagent. After 10 minutes of incubation, at 19°C and 200 RPM, relative luminescence units were recorded for each well from a top optic with a focal height of 11 mm and a gain of 3500 using a CLARIOstar plate reader (BMG Labtech).

After surface luminescence measurements, oocytes were washed for ≈30 seconds in modified Barth’s solution with gentle agitation. Luminescence measurement of permeabilized oocytes was performed subsequently using the kit Nano-Glo HiBiT Lytic Detection System (#N3040, Promega) by applying an equivalent protocol as described above. Measurements after lysis yield estimates of the total expression of HiBiT-tagged proteins.

Surface and total expression of a given construct was measured in several batches of oocytes expressing that construct. For each batch, luminescence values of 3 oocytes before and after lysis were measured and averaged. The resulting values were used to calculate averages over several batches of surface and total expressions.

#### Oocyte electrophysiology

Whole-cell currents were recorded with the two-electrode voltage clamp (TEVC) technique using a GeneClamp 500B Amplifier from Axon Instruments, a Digitizer Digidata 1440A from Axon Instruments and an eight-channel perfusion system with a manually operated controller from AutoMate Scientific. Some experiments where light stimulation was not needed were conducted with a HiClamp robot (MultiChannel System). Microelectrodes were filled with 3 M KCl and oocytes were perfused in the specified solutions. Bath solutions (composition in mM) were: ND96 (91 NaCl, 2 KCl, 1.8 CaCl_2_); ND96 0Ca (91 NaCl, 2 KCl); 94 K^+^ 100 Cl^-^ (94 KCl, 2 CaCl_2_); 94 Na^+^ 100 Cl^-^ (94 NaCl, 2 CaCl_2_); 94 K^+^ 10 Cl^-^ (4 KCl, 90 K Methanesulfonate, 2 CaCl_2_); 49 Ca^2+^ (49 CaCl_2_, 47 Glucose). All bath solutions had also 1 mM MgCl_2_ and 5 mM HEPES, and were adjusted to pH 7.4 with Tris or Citric acid, except for ND96 and ND96 0Ca where NaOH was used.

Currents were filtered at 3 kHz and sampled at 10 kHz.

Pulses of 505-nm light were applied with an OPTOLED LITE dual LED light source (Cairn Research Ltd; 5 A max LED current), coupled to a 1-mm optic fiber placed ∼3 cm above the oocyte. In that setup, the light, applied with a single optical fiber, can reach at best half of the oocyte surface and can penetrate a few microns of the dense cytoplasm of the oocyte. We estimate that 90% of the light is absorbed within a layer of <40 μm below the surface. This rough estimate is based on data from & Miledi (*48*) who found a 1/10 attenuation of 350-nm UV light at a distance <10 μm below the oocyte surface, and from Ash et al. (*49*) who showed that 500-nm light penetrates 4-time deeper in tissues than 350-nm light. Assuming that illumination penetrates 40 μm below the surface, we calculate that ∼5% of the intracellular volume of a 1.1-mm diameter oocyte is illuminated.

For oocytes injected with M3 mRNA, activation of the receptor was accomplished by perfusing 5 μM acetylcholine (#A6625, Sigma-Aldrich). In experiments with BAPTA_in_, each oocyte was injected with 50 nl of a 40 mM-BAPTA (#2786, Tocris) solution and was left incubating for at least 30 min before recording. This yields an overall internal BAPTA concentration of ∼3 mM in a stage VI oocyte of 1.1-mm diameter. In experiments where inhibitors were used the concentrations were as follows: 100 μM 2-APB (#1224, Tocris), 30 μM Ani9 (#6076, Tocris), 30 μM MONNA (#5770, Tocris) and 10 μM YM-254890 (#21910-1590, Tebu-BIO).

#### *Xenopus* tadpoles

Experiments on *Xenopus* tadpoles were carried out in accordance with the European Community Guide for Care and Use of Laboratory Animals and approved by the “Comité d′éthique en expérimentation de Bordeaux”, N° 33011005-A. Two independent batches of *Xenopus laevis* embryos were obtained and staged as previously described (*50, 51*). For each batch, at the two-cell stage, one cell of each embryo was injected with 250 pg lacZ (control) or 1 ng OLPVR1 mRNA.

#### *Xenopus* tadpoles behavior studies

Injected *Xenopus laevis* embryos were incubated in the dark in modified Barth’s solution for 3 to 4 days until tadpoles reach stage 39 to 44. Individual tadpoles were placed in 35 mm dishes under a trinocular microscope mounted with an INFINITY8-2M Camera (Teledyne Lumera) for video recording. Tadpoles were subjected to 2-200 s green-light pulses using either OPTOLED LITE dual LED light source (Cairn Research Ltd; 5 Amax LED current, 505 nm) or Alonefire X004, (China, 510∼530 nm).

#### Mammalian cells electrophysiology

HEK293T cells were maintained in DMEM supplemented with 10% FBS in 35-mm dishes. At 70–80% confluency, cells were transiently transfected using the calcium phosphate method (CaCl_2_ 2.5 M) with a total amount of 1.5 (OLPVR1) and 2.1 μg (TMEM16A, TMEM16B, SK1) of DNA, respectively, and seeded on 35-mm diameter plates.

Whole-cell patch-clamp recordings in HEK293T cells were performed 1-2 days after transfection. Recordings were performed in voltage-clamp mode using an Axopatch 200B (Molecular Devices) amplifier. The standard bath solution contained (in mM): 150 NaCl, 5 KCl, 10 HEPES (pH 7.4) and 2 CaCl_2_. The standard pipette solution contained (in mM): 155 KCl, 3 MgCl_2_, 10 HEPES (pH 7.3). Unless otherwise specified, it also contained 10 μM EGTA. EGTA in the pipette was used to chelate contaminant Ca^2+^. From specification sheets of salt (Sigma-Aldrich) we estimated contaminant Ca^2+^ to be ∼3 μM, and a predicted (Alex software (*52*)) free Ca^2+^ of 114 nM in 10 μM EGTA. Pulses of 505 nm-light were applied using a Thorlabs M505L4 LED mounted on a Zeiss Axiovert 200M microscope under a 40X objective.Signals were filtered at 10 kHz and sampled at 20 kHz.

#### Mammalian cells confocal fluorescence imaging

HEK293T cells were seeded in 25-mm poly-L-lysine-treated coverslips and transfected using Lipofectamine 2000 (ThermoFischer Scientific) with DNA for OLPVR1 fused to GFP at the C-terminal in pcDNA3.1, and pDsRed2-ER vector (Clontech, #632409) in a 6:1 ratio. After one day of expression, the cells were incubated for 30 min at 37ºC in standard bath solution with Hoescht 33342 (1/5000 dilution) and CellMask Deep Red (1/1000 dilution; #C10046 ThermoFischer Scientific). Hoescht 33342 was applied to stain the cell nucleus and CellMask Deep Red to stain the plasma membrane. pDsRed2-ER expression was used to label the ER. After the incubation, the cells were washed twice with standard bath solution and imaged immediately. Imaging was acquired under the objective Plan-Apochromat 63x/1.40 Oil DIC M27 of a Zeiss LSM 710 Laser Scanning Confocal Microscope equipped with the lasers DPSS 561-10 nm, Argon 488 nm, Diode 405-30 nm and Helium Neon 633 nm, as well as two descanned GaAsp detectors ([570-610 nm], [500-550 nm]) and a descanned PMT detector set to 415-485 nm or 642-740 nm. Composite images result from channel merging with ImageJ software.

#### Data analysis

For TEVC and patch-clamp recordings, data acquisition was performed using pClamp software (Molecular Devices). HEK cells data were analyzed with pClamp. Oocytes data were transferred to Microsoft Excel after undersampling to 10 or 100 Hz. Annotation, plotting, and analysis of the current recordings were automated by the use of Excel add-ins eeTevc and eeDataStat (*47*). Slow fluctuations of the baseline of the current signal were removed by interactive fitting of the baseline with a spline curve and subtraction of this fit from the signal. For luminescence experiments, unless otherwise noted, data points were blank-corrected and values shown represent the average of triplicates.

Statistical significance analysis was done using GraphPad Prism 8.

### Supplementary Text

#### Promoting surface expression of OLPVR1

Although exogenous membrane proteins have usually better surface expression in Xenopus oocytes than other systems (*53*), engineering surface-addressed OLPVR1 proved difficult (Fig. S6). Only one construct, the DOR=OLPVR1 fusion, displayed significant, though modest, surface expression, but its analysis was not pursued because (1) its photocurrents were too small (<5 nA) and (2) it functionally diverged from OLPVR1 as it did not induce Ca^2+^ release despite being highly enriched intracellularly.

The Organic Lake phycodnavirus genome where the OLPVR1 gene was found features another rhodopsin gene coding for OLPVR2 (*7*). In oocytes, OLPVR2 did not express at the surface and expressed only weakly (14-fold less than OLPVR1) intracellularly (Fig. S6). We were unable to record any electrophysiological activity associated with OLPVR2, native or with added signal sequences, in oocytes. Nor did we find that coexpression of OLPVR2 affected the properties of the photocurrents elicited by OLPVR1.

#### Aminoacid sequences of VRs used in this study

>OLPVR1

MDNIIMTAYISIFVQIITAIISVYGLFIPLNFKDIILREILILELIVQIIEFIFYIWLIITLQSINEDITYVRYFDWVLTTPVMLLTTVYFFEYMNSDDGIRKKEINDRDYVYLFYICLSNFFMLLIGYLGETKQINKMLTLFGGSFFLFLTFYLLYVKYTKENWMNYIVFYFMFLVWFLYGFAFMFPFSIKNQMYNILDIVSKNIYSIFIFIVILNQSYKLL

>VirChR1

MKDKELLLLTVKISLVVQIISGIVSSFGIFIKLAPKDYILRDILIIETVVQIVEAIFYVYIYLSLESLDNNVITSRRYFDWVITTPIMLISTILFMRYNIRILYDKSKNKQNRQNKPKSNLTTYNVIKNNKDTIIKIVIGNFIMLCFGYLGEVDILSKTISIPIGFIFFFYVFYLIWREFGSYTQQNQILFWVLFIVWGLYGVSACLPVLQKNICYNILDIISKNFYGLYIFYYIYQHRLVV

>TARA150

MVGGSLMRLTFYISIVIQFITGIVTFGGIFYKLPKQDRVLKDILGLETIVQLVEGLFYMYIISSLRYMRSNDITKRRYLDWVITTPMMLLSTIIYMEYENNKLTNKIVNTKEFINNNRKNITNMFLFNTLMLVSGFLGETGIMLKSMTIPIGFVFLFLAFNIMYKNYVGKADINKRIFMLMIIIWSMYGIAAMFPTILKNISYNILDIISKNFYGLYLFAKVRDIATRTT

>OLPVR2

MSDLIEYSFYLTYAFLMTTGTITFIEALRTKNESVRHILNLETCISVVAAFFYSNFIGKLEHINYEEINLNRYVDWAITTPIMLLVLVLAFRVNQTNKAMVKFSDFMIILGMNYGMLGTGYLGDIGVIHKTMGTVLGFLFFGGLFYKLNTLRTSNASNDLLYGAFFVLWALYGVFYQMEQLPRNVGYNVLDLFSKCFVGIYFWAFYAKIFT

**Fig. S1.**
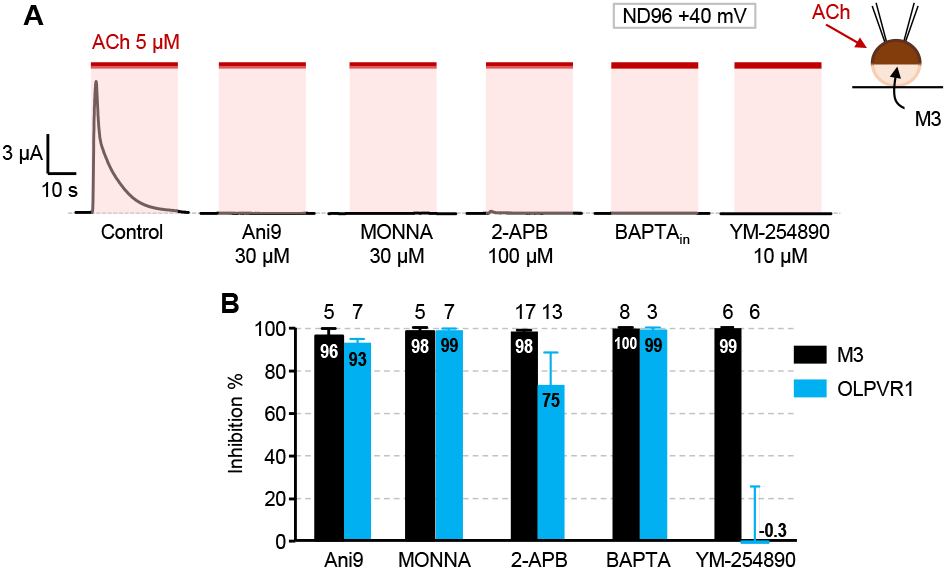
Comparison of the effects of inhibitors on OLPVR1 photocurrents and M3-mediated currents. (A) Representative currents elicited by 5 μM ACh in oocytes injected with 2.5 ng RNA coding for the Gq-coupled muscarinic M3 receptor. Bath solution = ND96, voltage = +40 mV. Oocytes were incubated in ND96 solution containing no inhibitor (Control), 30 μM Ani9, 30 μM MONNA, 100 μM 2-APB, or 10 μM YM-254890. Duration of incubation was 10’ for YM-254890, 60’ otherwise. BAPTA designates oocytes injected with 50 nl of a 40-mM BAPTA solution and left 60’ in ND96 solution prior to recording. The expected intracellular BAPTA concentration of BAPTA-injected oocytes is 3-4 mM for oocytes of diameter 1-1.1 mm. (B) Average inhibition of M3-induced currents in the conditions of panel A (M3; Black bars) and of OLPVR1 photocurrents measured at +40 mV in ND96 solution (OLPVR1; Blue bars). Values are specified in the bar. The numbers of oocytes tested in ND96 at 40 mV are indicated above the bars. Tests in other solutions and/or at different voltages gave similar results. The exact values for 2-APB inhibition of M3 and OLPVR1 currents were 98.1±0.47 (n=17) and 75.3±14.0 (n=13), respectively (Error bars, SEM).

**Fig. S2.**
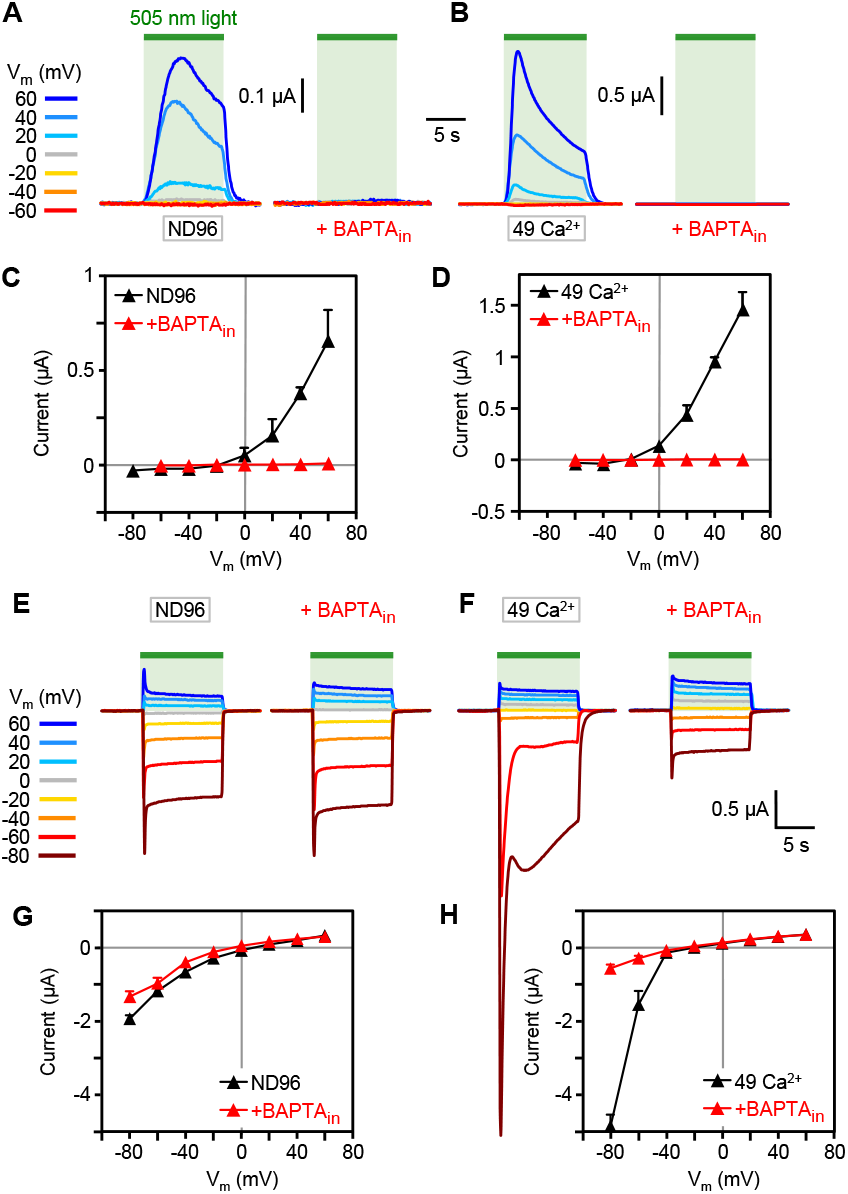
Unlike ChR2, OLPVR1 photocurrents are abrogated by intracellular injection of BAPTA. (A) Representative photocurrents from OLPVR1-expressing oocytes (7.5 ng RNA) with and without intracellular injection of BAPTA (BAPTA_in_) in ND96 bath solution (wich includes 1.8 mM Ca^2+^). (B) Same in 49 Ca^2+^ bath solution. (C) Average peak photocurrent vs voltage in ND96 solution with and without injected BAPTA. Each data point represents the average of 3-10 oocytes. (D) Same in 49 Ca^2+^ bath solution. Each data point represents the average of 4-5 oocytes. (E-F) Same as panels A and B, but with an oocyte expressing ChR2 (7.5 ng RNA). All traces are from the same oocyte. (G-H) Same as panels C and D, but with ChR2. Averages are from 4-5 oocytes. (Error bars, SEM).

**Fig. S3.**
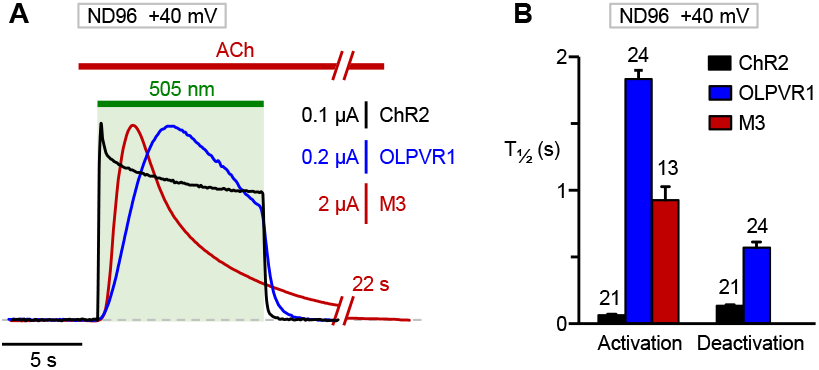
OLPVR1 photocurrents activates slowly upon illumination. (A) Representative currents elicited by illumination of oocytes expressing OLPVR1 (blue trace) and ChR2 (black trace), or application of 5 μM ACh to an oocyte expressing M3 receptors (brown trace; break in the trace represents 22 s of omitted data). Superimposed traces were scaled vertically to the same size, vertical scale bars for each trace are indicated. Bath solution = ND96 solution, voltage = +40 mV. (B) Average half-times of current activation upon light or ACh application, and deactivation upon light switch-off. Conditions were as in panel A. Numbers above bars represent the numbers of oocytes analyzed. The values for M3 were calculated as the time interval between the onset of the increase in current after ACh application and the time where current reached 50% of its peak value. They therefore do not include the latency inherent to bath application of agonists to oocytes. The time resolution of the recordings is ∼50 ms so that the measured activation and deactivation T_½_ of ChR2 of 65±6 and 135±9 ms, respectively, are upper estimates of the actual values.

**Fig. S4.**
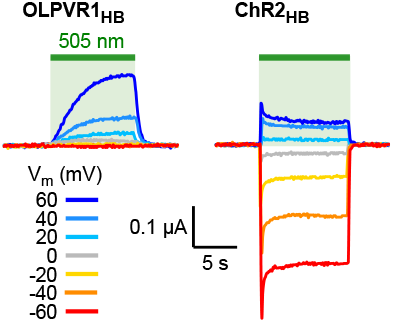
HiBit tag does not modify the function of rhodopsins. Representative photocurrents of HiBit-tagged OLPVR1 (OLPVR1_HB_; 30 ng RNA/oocyte) and ChR2 (ChR2_HB_; 7.5 ngRNA/oocyte) in ND96 solution.

**Fig. S5.**
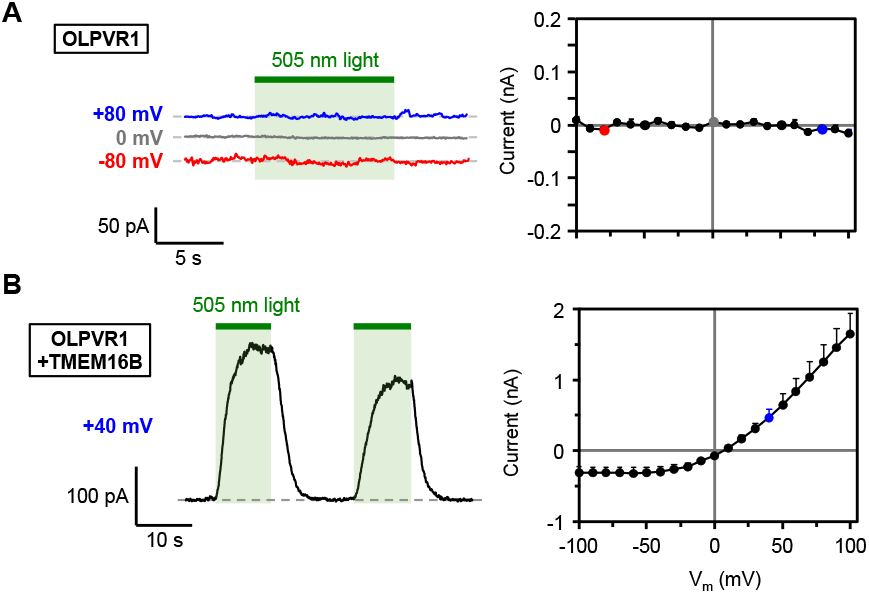
OLPVR1 activates surface Ca^2+^-activated TMEM16B channels in mammalian cells through release of intracellular calcium. Data were obtained from HEK293T cells transfected with OLPVR1 alone (A) or with OLPVR1 and TMEM16B (B). Left panels : Representative current responses to illumination (green bars, 505 nm, 10 s). Cells were held at the indicated voltages. Right panels: Light-induced current (peak current elicited by first illumination – current before illumination) vs. voltage. Currents were elicited by voltage ramps from -100 to +100 mV (400 ms duration) repeated every second. Averages were calculated from 8 cells in each conditions (Error bars, SEM). The bath solution contained (in mM): 150 NaCl, 5 KCl, 2 CaCl_2_, and 10 HEPES (pH 7.4). The pipette solution contained (in mM): 155 KCl, 3 MgCl_2_, 10 HEPES (pH 7.3) and 10 μM EGTA.

**Fig. S6.**
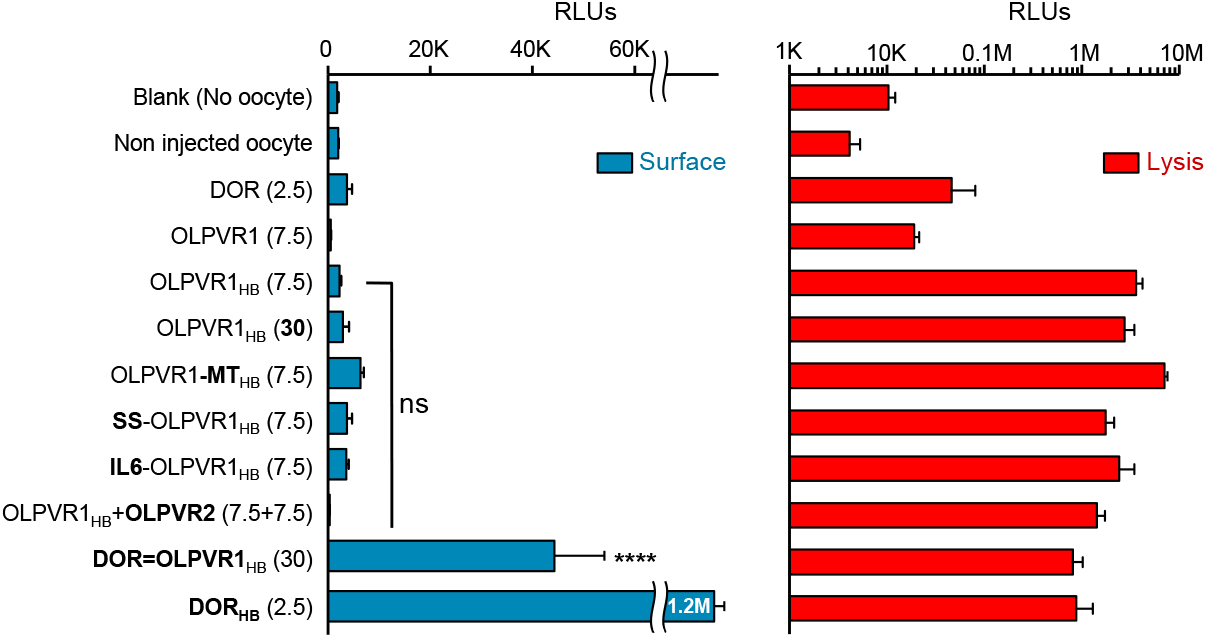
Surface expression of various OLPVR1 constructs. Mean luminescence (left: surface values, right: values after lysis) recorded in control conditions (no oocyte and non-injected oocytes) or from RNA-injected oocytes. Luminescence values were not blank corrected and averages were calculated from all oocytes tested (n=3-114). Amounts of RNA coding for specified proteins, in ng per oocyte, are indicated in parenthesis. HB suffix indicated the presence of a HiBiT tag at the N-terminus. Oocytes were maintained in ND96 solution supplemented with 1 μM all-trans-retinal and tested 24-72h after injection. Increasing the amount of OLPVR1 RNA injected per oocyte from 7.5 to 30 ng did not affect surface expression. Modifying the protein by inserting signal sequences did not improve surface expression (MT, Golgi export trafficking signal of Kir2.1 potassium channel; SS, Signal sequence of human nicotinic acethylcoline alpha 7 receptor subunit; IL6, Interleukine 6 secretion signal sequence). Coexpressing OLPVR1 with viral rhodopsin OLPVR2 did not improve surface expression. However, the fusion DOR=OLPVR1 (30 ng) had a surface expression 15-fold higher than OLPVR1 (30 ng), but still 25-fold lower than DOR alone (DOR, δ-opioid receptor). **** P<0.0001. ns P>0.9999, not significant. One-way ANOVA followed by Dunnett’s multiple comparison test against the control OLPVR1 (7.5 ng).

### Movie S1

#### OLPVR1-injected tadpole swims in response to green light

File: SV1_OLPVR1_1ng_day3_GreenLight_Swimming.mp4

*OLPVR1 Tadpole 4 days after injection of 1 ng OLPVR1 RNA*.

*Test: 1 application (∼30 s) of green light (wavelength?). Result = swimming*.

### Movie S2

#### OLPVR1-injected tadpole flicks tail in response to green light

File: SV2_OLPVR1_1ng_day4_GreenLight_TailFlick.mp4

*OLPVR1 Tadpole 4 days after injection of 1 ng OLPVR1 RNA*.

*Test: 3 applications (∼2 s) of green light (510-530 nm). Result = tail flicking*.

### Movie S3

#### OLPVR1-injected tadpole does not respond to red light

File: SV3_OLPVR1_1ng_day4_RedLight.mp4

*OLPVR1 Tadpole 4 days after injection of 1 ng OLPVR1 RNA*.

*Test: 4 applications (∼5 s) of red light (600-720 nm). Result = no response*.

### Movie S4

#### LacZ-injected tadpole does not respond to green light

File: SV4_LacZ_1ng_day3_GreenLight.mp4

*OLPVR1 Tadpole 4 days after injection of 1 ng LacZ RNA*.

*Test: 1 prolonged application (∼35 s) of green light (510-530 nm). Result = no response*.

